# LEARNING AND INTERPRETING THE GENE REGULATORY GRAMMAR IN A DEEP LEARNING FRAMEWORK

**DOI:** 10.1101/864058

**Authors:** Ling Chen, John A. Capra

**Affiliations:** Department of Biological Sciences, Vanderbilt University, Nashville, TN, 37235, USA; Vanderbilt Genetics Institute and Department of Biomedical Informatics, Vanderbilt University Medical Center, Nashville, TN 37235, USA; Department of Computer Science, Vanderbilt University, Nashville, TN, 37235, USA

## Abstract

Deep neural networks (DNNs) have achieved state-of-the-art performance in identifying gene regulatory sequences, but they have provided limited insight into the biology of regulatory elements due to the difficulty of interpreting the complex features they learn. Several models of how combinatorial binding of transcription factors, i.e. the regulatory grammar, drives enhancer activity have been proposed, ranging from the flexible TF billboard model to the stringent enhanceosome model. However, there is limited knowledge of the prevalence of these (or other) sequence architectures across enhancers. Here we perform several hypothesis-driven analyses to explore the ability of DNNs to learn the regulatory grammar of enhancers. We created a synthetic dataset based on existing hypotheses about combinatorial transcription factor binding site (TFBS) patterns, including homotypic clusters, heterotypic clusters, and enhanceosomes, from real TF binding motifs from diverse TF families. We then trained deep residual neural networks (ResNets) to model the sequences under a range of scenarios that reflect real-world multi-label regulatory sequence prediction tasks. We developed a gradient-based unsupervised clustering method to extract the patterns learned by the ResNet models. We demonstrated that simulated regulatory grammars are best learned in the penultimate layer of the ResNets, and the proposed method can accurately retrieve the regulatory grammar even when there is heterogeneity in the enhancer categories and a large fraction of TFBS outside of the regulatory grammar. However, we also identify common scenarios where ResNets fail to learn simulated regulatory grammars. Our results provide a framework for interpreting the regulatory rules learned by ResNets, and they demonstrate that the ability and efficiency of ResNets in learning the regulatory grammar highly depends on the nature of the prediction task.

## Introduction

Enhancers are genomic regions distal to promoters that regulate the dynamic spatiotemporal patterns of gene expression required for the proper differentiation and development of multi-cellular organisms (Kundaje et al., 2015; Shlyueva et al., 2014; Villar et al., 2015). As a result of their essential role, mutations that disrupt proper enhancer activity can lead to diseases. Indeed, the majority of genetic variants associated with complex disease in genome-wide association studies (GWAS) are non-protein coding, and thought to influence disease by disrupting proper gene expression levels (Brazel and Vernimmen, 2016; Corradin and Scacheri, 2014; Maurano et al., 2012).

Enhancers function through the coordinated binding of transcription factors (TFs). Recent advances in high-throughput sequencing techniques have greatly deepened our knowledge of TF binding specifies (Bernstein et al., 2012; Jolma et al., 2013; Lambert et al., 2018). However, identifying consensus TF binding motifs are not sufficient for inferring TF binding. As shown in many ChIP-seq studies, TFs only bind to a small fraction of all motif occurrences in the genome, and some binding sites do not contain the consensus TF binding motif, indicating a necessity for additional features (Wang et al., 2012). Indeed, many additional features have been suggested to play a role in determining *in vivo* TF binding, such as heterogeneity of a TF’s binding motif (Samuel Levy and Hannenhalli, 2002), local DNA properties (Dror et al., 2015), broader sequence context and interposition dependence (Mathelier and Wasserman, 2013), cooperative binding of the TF with its partners (Jolma et al., 2015; Liu et al., 2015; Wang et al., 2006; Yáñez-Cuna et al., 2013), and condition-specific chromatin context (Heintzman et al., 2009; Kumar and Bucher, 2016; Wang et al., 2006). While both genomic and epigenomic features are important in determining the *in vivo* occupancy of a TF, recent studies have suggested that the epigenome can be accurately predicted from genomic context (Arvey et al., 2012; Benveniste et al., 2014; Dror et al., 2015; Whitaker et al., 2015), supporting the fundamental role of sequence in dictating the binding of TFs (Li and Ovcharenko, 2015; Prescott et al., 2015; Ritter et al., 2010; Schmidt et al., 2010; Wilson et al., 2008). Therefore, it is critical to understand the sequence patterns underlying enhancer regulatory functions and build sufficiently sophisticated models of enhancer sequence architecture.

Combinatorial binding of TFs, i.e., the regulatory “grammar” that combines TF “words”, is thought to be essential in determining *in vivo* condition-specific binding (Arvey et al., 2012; S Levy and Hannenhalli, 2002; Mathelier and Wasserman, 2013; Sharmin et al., 2015). However, how enhancers integrate multiple TF inputs to direct precise patterns of gene expression is not well understood. Most enhancers likely fall on a spectrum represented by two extreme models of enhancer architecture: the *enhanceosome model* and the *billboard model* (Long et al., 2016; Slattery et al., 2014). The enhanceosome model proposes that enhancer activity is dependent on the cooperative assembly of a set of TFs at enhancers. The cooperative assembly of an enhanceosome is based on physical protein-protein interactions and highly constrained patterns of TF-DNA binding sites. The enhanceosome model does not tolerate shifts in the spacing, orientation, or ordering of the binding site, which can disrupt protein-protein interactions and cooperativity. This model likely presents an extreme example because only very few enhancers are found under such stringent constraints (Crocker et al., 2008; Erives and Levine, 2004; Papatsenko and Levine, 2007; Swanson et al., 2011, 2010). However, many examples of less extreme spatial constraints on paired TF-TF co-association and binding-site combinations are found in genome-wide ChIP sequencing studies (Cheng et al., 2013; Kazemian et al., 2013; Sorge et al., 2012) and *in vitro* consecutive affinity-purification systematic evolution of ligands by exponential enrichment (CAP-SELEX) studies. On the other end of the spectrum is the billboard model, also known as the information display model (Arnosti and Kulkarni, 2005; Kulkarni and Arnosti, 2003), which hypothesizes that instead of functioning as a cooperative unit, enhancers work as an ensemble of separate elements that independently affect gene expression. That is, the positioning of binding sites within an enhancer is not subject to strict spacing, orientation, or ordering rules. The TFs at billboard enhancers work together to direct precise patterns of gene expression, but their function does not strongly depend on each other. For instance, the loss of a TF binding may lead to change in the target gene expression, but will not cause the complete collapse of enhancer function. The actual mechanisms by which multiple TFs assemble on enhancers are likely a mixture of the two models. Indeed, a massively parallel reporter assay (MPRA) of synthetic regulatory sequences suggested that while certain transcription factors act as direct drivers of gene expression in homotypic clusters of binding sites, independent of spacing between sites, others function only synergistically (Smith et al., 2013).

In recent years, deep neural networks (DNNs) have achieved state-of-art prediction =accuracies for many tasks in regulatory genomics, such as predicting splicing activity (Leung et al., 2014; Xiong et al., 2015), specificities of DNA- and RNA-binding proteins (Alipanahi et al., 2015), transcription factor binding sites (TFBS) (Quang and Xie, 2019, 2015; Zhou and Troyanskaya, 2015), epigenetic marks (Kelley et al., 2016; Quang and Xie, 2015; Zhou and Troyanskaya, 2015), enhancer activity (Min et al., 2016; Yang et al., 2017) and enhancer-promoter interactions (Singh and Yang, 2016). However, in spite of their superior performance, little biological knowledge or mechanistic understanding has been gained from DNN models. In computer vision, the interpretation of DNNs trained on image classification tasks demonstrate that high-level neurons often learn increasingly complex patterns building on those learned by lower level neurons (Olah et al., 2018, 2017; Shrikumar et al., 2017; Springenberg et al., 2014; Yosinski et al., 2015; Zeiler and Fergus, 2014; Zeiler et al., 2010). DNNs trained on DNA sequences might behave similarly, with neurons in low levels learning building blocks of the regulatory grammar, short TF motifs, and those in higher levels learning the regulatory grammar itself, the combinatorial binding rules of TFs (Chen et al., 2018; Kelley et al., 2016; Quang and Xie, 2016).

The majority of DNNs trained with genomic sequences use a convolution layer as a first layer and then stack convolution or recurrent layers on top of it. A common approach to interpret the features learned by such DNNs is to convert the first convolution layer neurons to position weight matrices by counting nucleotide occurrences in the set of input sequences that activate the neurons (Alipanahi et al., 2015; Chen et al., 2018; Kelley et al., 2016). With the development of more advanced DNN visualization and interpretation techniques in computer vision, many other DNN interpretation methods emerged, such as occlusion (Zeiler and Fergus, 2014), saliency maps (Simonyan et al., 2013), guided propagation (Zeiler and Fergus, 2014), gradient ascent (Yosinski et al., 2015). Some of these techniques have been applied to visualize features learned by DNNs trained with genomic sequences. For instance, a gradient based approach, DeepLIFT, identified relevant transcription factor motifs in the input sequences learned by a convolutional neural network (Shrikumar et al., 2017). Saliency maps, gradient ascent and temporal output scores have been used to visualize the sequence features learned by a DNN model for TFBS classification and found informative TF motifs (Lanchantin et al., 2017). These studies demonstrate the power of DNNs in recognizing the TF motifs in the input sequences. However,these studies focused only on the interpretation of the output layer in models for predicting TFBS. Enhancers can be much more complex than individual TFBS as they contain multiple binding sites in range of combinations and organizations. It is also unclear whether the intermediate layers of DNNs have the capability of learning rules of combinatorial TF binding from regulatory regions with many TFs, such as enhancers.

Another substantial challenge in the development of methods to interpret DNNs applied to regulatory sequences is our lack of knowledge of the combinatorial rules governing enhancer function in different cell types. Beyond a few foundational examples used to propose possible enhancer architectures, the constraints and interactions that drive enhancer function are largely unknown. Thus, it is difficult to determine if a pattern learned by a neuron is “correct” or biologically relevant. The generation of synthetic DNA sequences that reflect different constraints on regulatory element function has promise to help address these challenges and enable evaluation of the ability of DNNs to learn different regulatory architectures and of algorithms for reconstructing these patterns from the trained networks. Indeed, DeepResolve was recently proposed to interpret the combinatorial logic from intermediate layers of DNNs, and the ability of the neural network to learn the AND, OR, NOT and XOR of two short sequence patterns was demonstrated in a synthetic dataset (Liu and Gifford, n.d.). However, these simulated combinatorial logics and sequence patterns were not biologically motivated and were simpler than most proposed enhancer architectures.

Here, we develop a biologically motivated framework for simulating enhancer sequences with different regulatory architectures, including homotypic clusters, heterotypic clusters, and enhanceosomes, based on real TF motifs from diverse TF families. We then apply a state-of-the-art variant of deep neural networks, residual neural network (ResNet) algorithms, to classify these sequences and use this framework to investigate whether the intermediate layers the networks learn the complex combinatorial TF architectures present in the simulated regulatory grammars. In particular, we developed an unsupervised method for assigning transcription factor binding sites to grammars based on the gradients assigned to their nucleotides by intermediate layers of the neural network. We evaluate the efficiency in extracting simulated regulatory grammars under a range of scenarios that mimic real-world multi-label regulatory sequence prediction tasks, considering possible heterogeneity in the output enhancer categories and fraction of TFBS not in the regulatory grammar. We demonstrate that ResNet can accurately model simulated regulatory grammars in many multi-label enhancer prediction tasks, even when there is heterogeneity in the output categories or a large fraction of TFBS outside of regulatory grammar. We also identified scenarios where the ResNet fails to learn an accurate representation of the regulatory grammar, including using inappropriate control sequences as negative training examples, considering output categories differing in multiple sequence features, and having an overwhelming amount of TFBS outside of the regulatory grammar. In summary, our work makes three main contributions: i) We provide a flexible tool for simulating regulatory sequences based on biologically driven hypotheses about regulatory grammars. ii) We develop and evaluate an algorithm for interpreting the regulatory grammar from the intermediate layers of DNNs trained on enhancer DNA sequences. iii) We demonstrate that the ability of DNNs to learn interpretable regulatory grammars is highly dependent on the design of the prediction task.

## Results

To evaluate the performance of ResNets on modeling the regulatory grammar, we performed a simulation analysis (Figure 1a). We designed a set of 12 biologically motivated regulatory grammars consisting of TFs from diverse families. These include five homotypic clusters of the same TF, five heterotypic clusters of different TFs, and two enhanceosomes of different TFs with requirements on the spacing and orientation of their binding sites. Motivated by the fact that enhancers active in a given cellular context likely consist of multiple types with different grammars, we designed twelve “classes” of regulatory sequences. Each class contains a different set of regulatory grammars, but the grammars can occur within multiple classes, and TFs can occur within multiple grammars. Then, using these classes, we simulated 30,000 enhancer sequences which each contain a sequence that matches the pattern defined by one of the classes (Methods).

**Figure 1.**
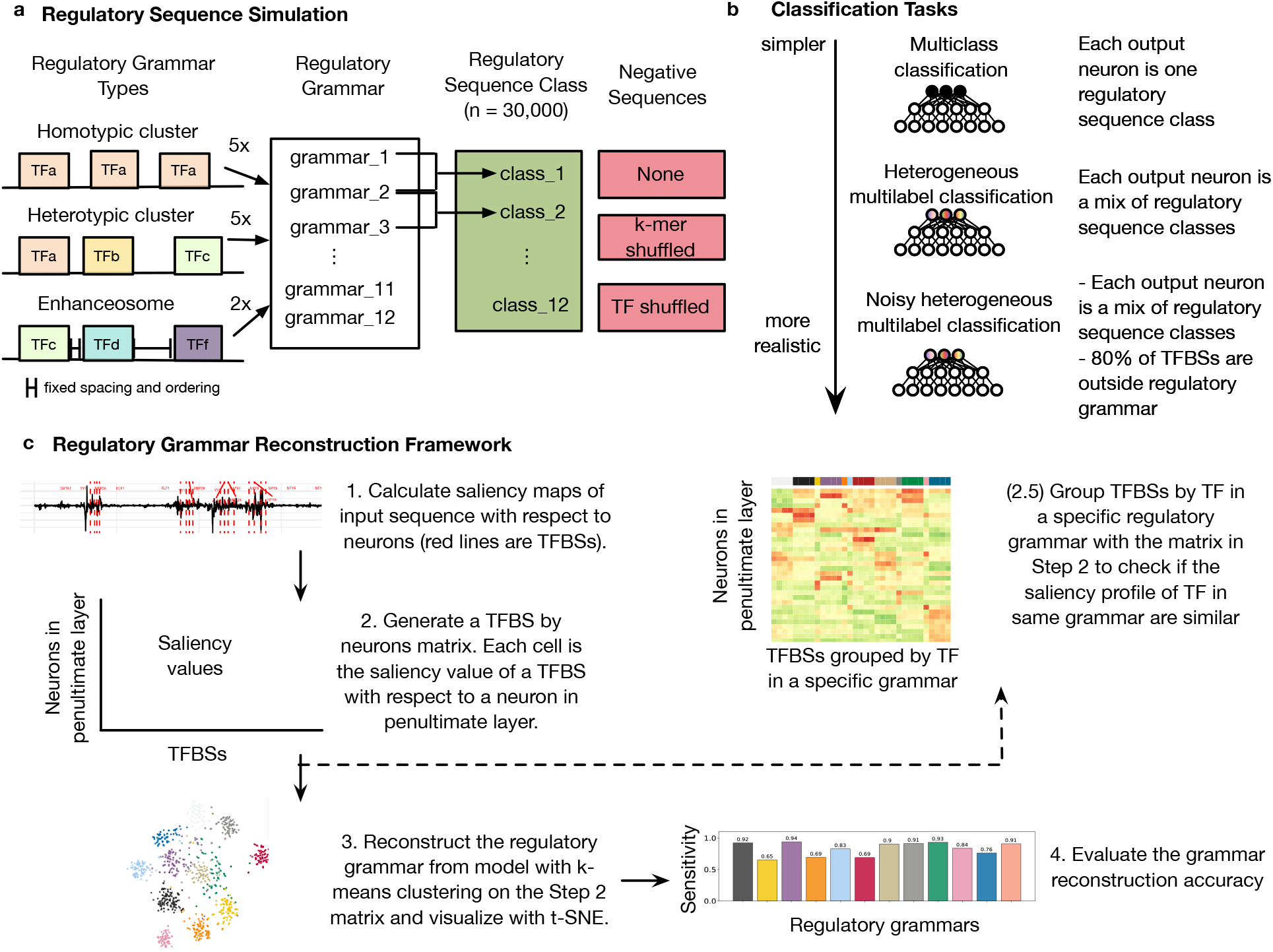
Pipeline for analyzing regulatory grammar learned by ResNet models trained on simulated regulatory sequences. (a) Regulatory sequence and negative sequences simulation. We designed twelve regulatory grammars, including five homotypic clusters, five heterotypic clusters, and two enhanceosomes as prototypes for simulated regulatory sequences. Then, to reflect that regulatory regions active in a cellular context may have multiple grammars, we defined twelve regulatory sequence classes, each with two different grammars. Finally, we generated two sets of negative sequences: k-mer shuffled and TF shuffled versions of the simulated positive sequences. (b) Classification tasks. ResNets are trained on simulated regulatory sequences and the negative sets in three increasingly realistic scenarios. (c) Regulatory grammar reconstruction framework.

Our goal is to evaluate the ability of ResNets to learn regulatory grammars and the ability of our proposed framework to reconstruct and visualize these grammars. Using sequences generated from the simulated regulatory grammars, we trained several models corresponding to different classification scenarios found in real-world regulatory sequence prediction tasks (Figure 1b). First, we trained a ResNet on a multi-class classification task using sequences from each of the regulatory classes and TF-shuffled negative sequences. Then, we investigated how well the approach performed when trained in more challenging situations, including no negative training sequences, k-mer matched negatives, heterogeneity in the output categories, and large fractions of TFBSs outside of regulatory grammars in the input sequences.

### ResNet trained on simulated regulatory sequences and TF-shuffled negatives accurately captures simulated regulatory grammars

To explore whether the ResNet model can learn the regulatory grammar, we started with a multi-class classification task based on simulated regulatory sequences from 12 classes and TF-shuffled negative sequences (Methods; Supplementary Table 1 and 2). We trained a classifier to predict the class of the sequence, either not a regulatory sequence or member of one of the regulatory sequence classes. By constructing the prediction task with TF matched negative sequences, the neural network is forced not only to learn the individual TF motifs, but also learn the combinatorial patterns between the TFs.

The ResNet model accurately predicts the class label of input DNA sequences with near perfect performance: average area under the ROC curve (auROC) of 0.999 and average area under the precision-recall curve (auPR) of 0.982. We then analyzed what features were learned by calculating saliency maps (Methods) of input sequences with respect to each neuron in the penultimate layer (the dense layer immediately before the output layer). We found that neurons in the penultimate layer detect the location of the simulated TFBS. For instance, when we compute the saliency map of a class 6 simulated regulatory sequence with respect to neuron 1 in the penultimate layer, the TFBS have higher saliency value compared to other locations in the sequence, indicating the higher importance of those nucleotides to the activation of neuron 1 (Figure 2a).

**Figure 2.**
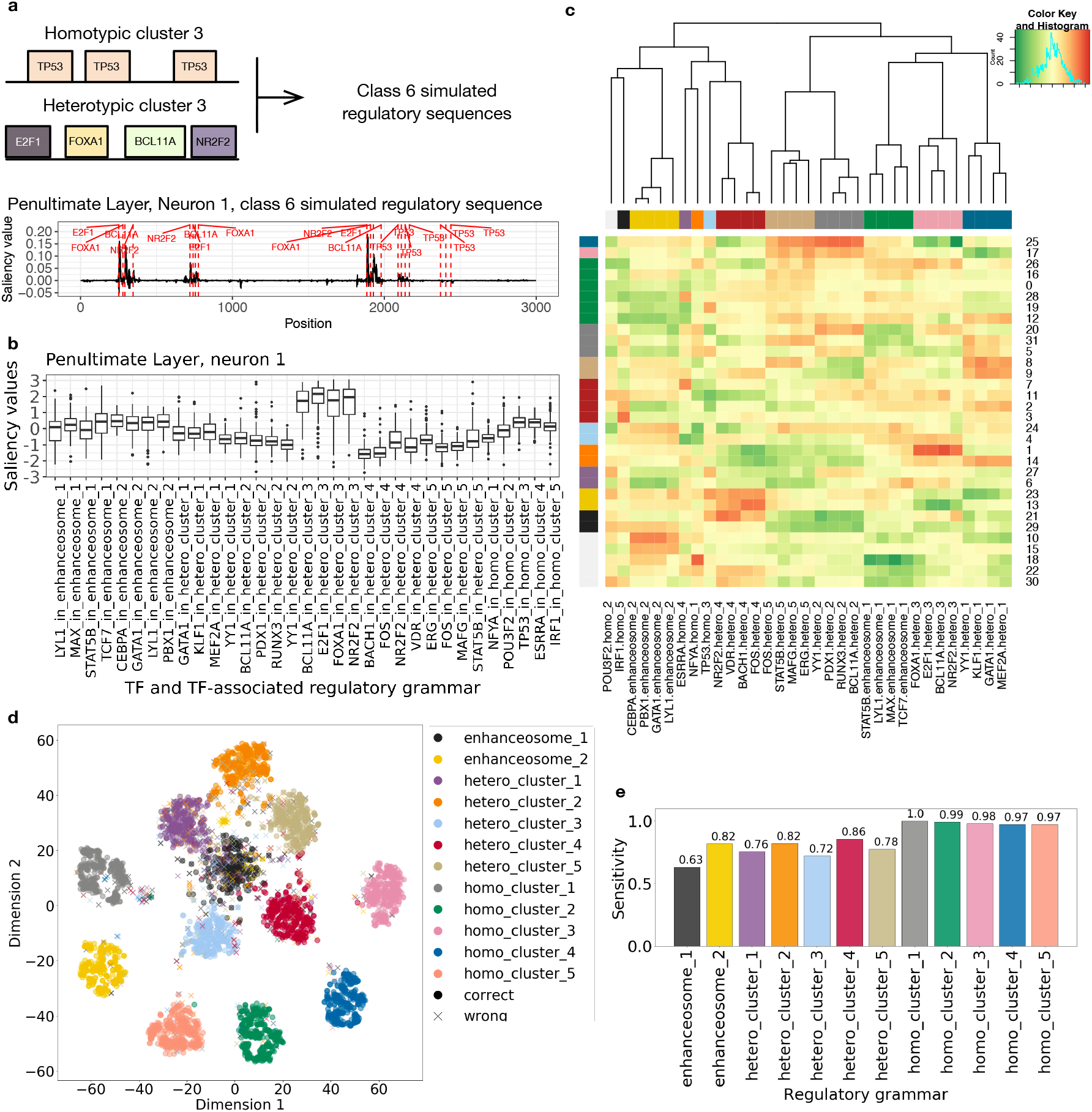
ResNet trained on simulated regulatory sequences and TF-shuffled negatives accurately models the regulatory grammar. (a) Illustration of the saliency map for simulated regulatory sequence from class 6. Class 6 sequences harbor instances of homotypic cluster 3 and heterotypic cluster 3. The saliency map shown is computed with respect to neuron 1 in the penultimate layer. The red dashed lines showed simulated TFBSs in their respective regulatory grammar. (b) The saliency values of the binding sites of each TF in a specific regulatory grammar with respect to neuron 1 in the penultimate layer. (c) Heatmap of the median saliency value of the binding sites of each TF in a specific regulatory grammar (x axis) across neurons of the penultimate layer (y axis). The order of x and y axis labels are determined by hierarchical clustering. The color bars on the side indicate the group label assigned by hierarchical clustering. (d) Actual labels of simulated regulatory grammar of the TFBS overlaid on t-SNE visualization of TFBS saliency values across neurons. Correct predictions of the regulatory grammar for a TF is represented by a dot, that is the predicted label agree with the actual label. Incorrect predictions of the regulatory grammar of a TF are indicated by crosses. (e) The sensitivity (TP/(TP+FN)) of the regulatory grammar predictions.

Next, we visualized the features learned by neuron 1 of the penultimate layer by plotting the mean saliency value of a 10 bp window from the start of each TF binding site using 240 sequences from all simulated regulatory sequence classes (Figure 2b). For example, the TFBS from heterotypic cluster 3 have elevated gradients compared to TFBS from other simulated regulatory grammars. This suggests that neuron 1 of the penultimate layer detects TFBS from heterotypic cluster 3. We then took the median saliency values of TFBSs in a specific regulatory-grammar and generated a matrix with rows of neurons and columns of each TF. As shown in Figure 2b, although TFBSs from heterotypic cluster 3 have elevated saliency values, those values are not always at the same level (E2F1 binding sites have higher median saliency value than the other three TFs). Therefore we scaled the matrix column-wise to identify which TF is most learned by which neuron. We plotted the scaled matrix as a heatmap with hierarchical clustering (Method; Figure 2c). We found that: (i) TFBSs from the same regulatory grammar have elevated gradients together and therefore are clustered; (ii) neurons of the penultimate layer can “multitask”, that is, one neuron can detect one or more regulatory grammars. For instance, neuron 25 in the penultimate layer learned both heterotypic cluster 2 and 5. This suggests that the penultimate layer captured the simulated regulatory grammars.

In order to evaluate how well the regulatory grammar can be reconstructed from the penultimate layer, we performed unsupervised clustering of TFBS based on their saliency values with respect to the neurons in the penultimate layer. More specifically, we performed a k-means clustering (k=12) of TFBSs from 240 sequences using their saliency values with respect to each neuron of the penultimate layer and visualized it with t-SNE (Figure 2d). Each TFBS has a predicted clustering label that is assigned by the k-means clustering algorithm and a true regulatory grammar. We first used majority voting to determine the predicted regulatory grammar for a cluster. For instance, the majority of cluster 1 is from heterotypic cluster 1, so we assign heterotypic cluster 1 as the predicted regulatory grammar for all TFBS in cluster 1. We then calculate the accuracy and sensitivity of the regulatory grammar reconstruction by comparing the predicted regulatory grammar and the true regulatory grammar. On average, 85.1% of TFBS are correctly classified (Figure 2e), and homotypic clusters are generally learned better (sensitivity > 0.97) than heterotypic clusters and enhanceosomes.

The same analysis approach can be applied to any layer of the neural network. We found that the neural network built up its representation of the regulatory grammar by first learning the individual TF motifs in the lower level neurons and gradually grouping TF motifs in the same regulatory grammar together (Supplementary Figure 3).

Taken together, these results demonstrate that ResNet models can largely capture simulated regulatory grammars if trained to perform a multi-class prediction with TF-shuffled negatives,and that our unsupervised clustering method based on saliency maps is able to reconstruct the regulatory grammar.

### Regulatory grammar can be learned by the ResNet model without TF-shuffled negatives

Although the ResNet model demonstrated the ability to capture the simulated regulatory grammars when trained against TF-shuffled negatives, we cannot construct perfect TF-shuffled negatives in the real-world, because the true TFs are not known. Indeed, in many applications, only the positive regulatory sequences (Quang and Xie, 2016; Zhou et al., 2018; Zhou and Troyanskaya, 2015) or k-mer shuffled negatives are used for training machine learning models. Therefore, we tested whether the ResNet model can learn the simulated regulatory grammar if trained with no negatives or k-mer shuffled negatives.

We trained five models for multi-class classification against: no negatives, 1-mer shuffled negatives, 4-mer shuffled negatives, 8-mer shuffled negatives, and 12-mer shuffled negatives. Then, we evaluate their performance at predicting simulated regulatory sequences. The model trained with 8-mer shuffled negatives achieved the highest accuracy at distinguishing TF-shuffled negatives from simulated regulatory sequences (average auROC 0.998, auPR 0.957, Figure 3a).

**Figure 3.**
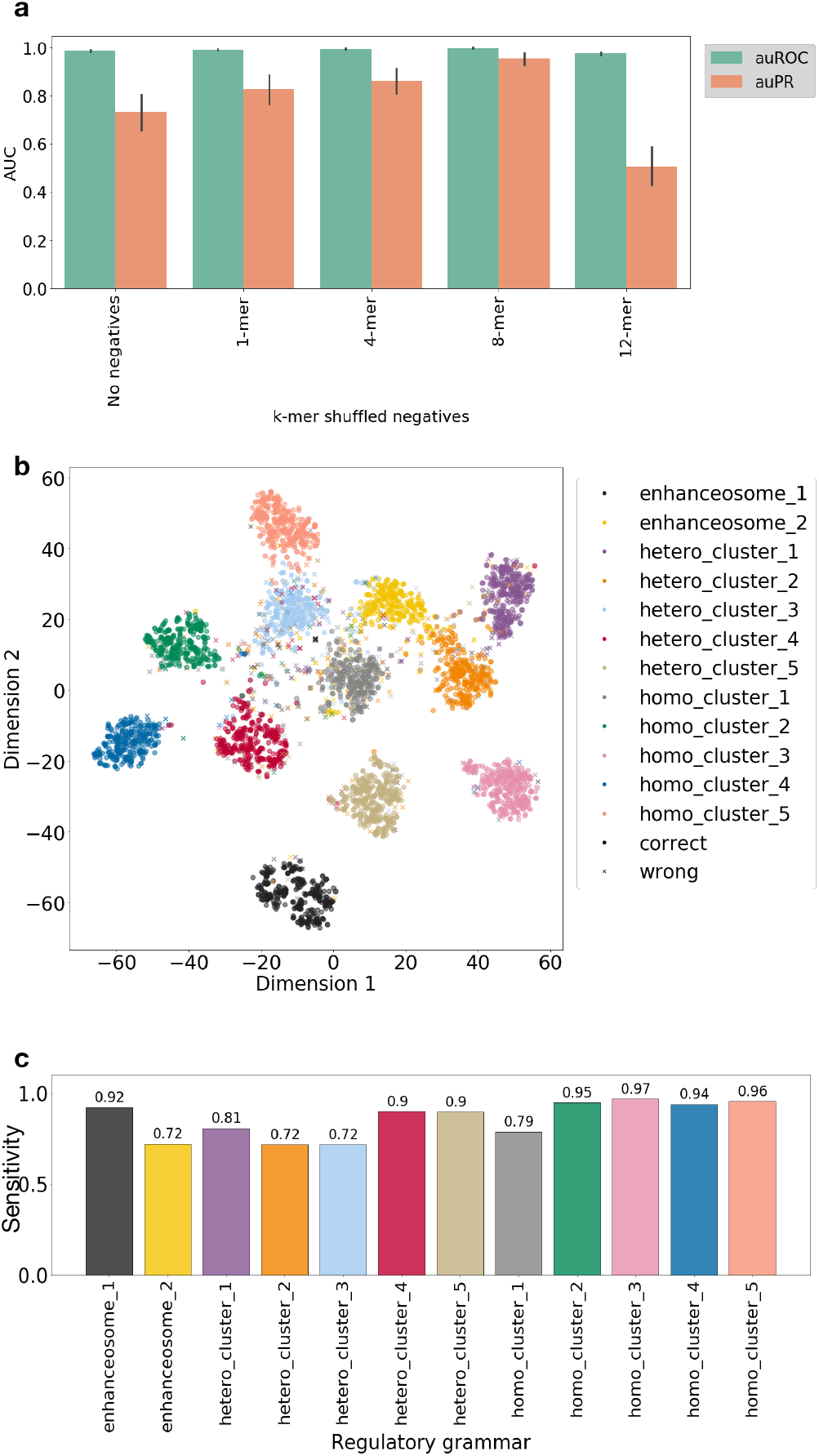
ResNet trained on simulated regulatory sequences against 8-mer shuffled negatives accurately models the regulatory grammar. (a) The performance of five different ResNet models trained on simulated regulatory sequences against different k-mer shuffled negatives at predicting the regulatory class of the simulated regulatory sequences vs. TFs-shuffled negatives test dataset. (b) Actual labels of simulated regulatory grammar of the TFBS overlaid on t-SNE visualization of TFBS saliency values across neurons. (c) The sensitivity of predicted labels in (b) of the ResNet model trained on the simulated regulatory sequences against 8-mer shuffled negatives.

To further explore the regulatory grammar learned by the ResNet model trained against 8-mer shuffled negatives, we calculated saliency maps over a set of input sequences (n=240) from each class of simulated regulatory sequences with respect to neurons in the penultimate layer. We performed hierarchical clustering on the median saliency values for the binding sites for each TF in a specific regulatory grammar as we did in the previous results section (Supplementary Figure 4). We found that TFBS from the same regulatory grammar were grouped together. Next, we performed k-means clustering (k=12) of the TFBS from the 240 sequences and overlaid the clustering label on the tSNE visualization (Figure 3b). We calculated the accuracy of predicted regulatory grammar for each TF. The average grammar reconstruction accuracy of this model is on par with the model trained against TF-shuffled negatives (85.3% vs. 85.1%).

These results suggest that the model trained against 8-mer shuffled negatives can learn a good representation of the regulatory grammar and therefore 8-mer shuffled negatives can be used as a substitute for TF-shuffled negatives in practice.

### Regulatory grammar can be learned by the ResNet model in the presence of heterogeneity in the regulatory sequences

A common task in regulatory sequence prediction is to predict sequences that exert a certain set of functions, e.g., activity in different cellular contexts. It is likely that sequences with a heterogeneous set of grammars are active in each cellular context.

To mimic this type of heterogeneity, we performed a heterogenous multi-label classification by pooling a number of simulated regulatory classes together as one heterogeneous class to generate five heterogeneous classes (Method; Fig 1b, Supplementary Table 4). We also allowed one regulatory class to be used in several heterogeneous classes. For example, in our simulation, regulatory sequences in heterogenous class 1 consist of regulatory class 1, 3, and 5. Regulatory class 1 sequences also belong to heterogenous class 5, and regulatory class 5 sequences also belong to heterogenous class 4. This multi-function of a regulatory sequence class is often observed in real-word regulatory sequences as many enhancers are active in more than one cellular context.

We trained the ResNet model against k-mer shuffled negatives (k=1, 4, 8, 12). Again, the model trained against 8-mer shuffled negatives performed the best when evaluated against the TF-shuffled negatives (average auROC 0.99, auPR 0.93, Supplementary Figure 5a). We performed hierarchical clustering (Supplementary Figure 5b) and unsupervised clustering (Figure 4a, b) as we did in the previous sections. The model trained to predict the heterogenous classes can still learn the majority of the regulatory grammars. The average accuracy of reconstructing regulatory grammar in this setting is 89.2%, which is similar to that of the multi-class classifications against TF-shuffled negatives (85.1%) and against k-mer shuffled negatives (85.3%).

**Figure 4.**
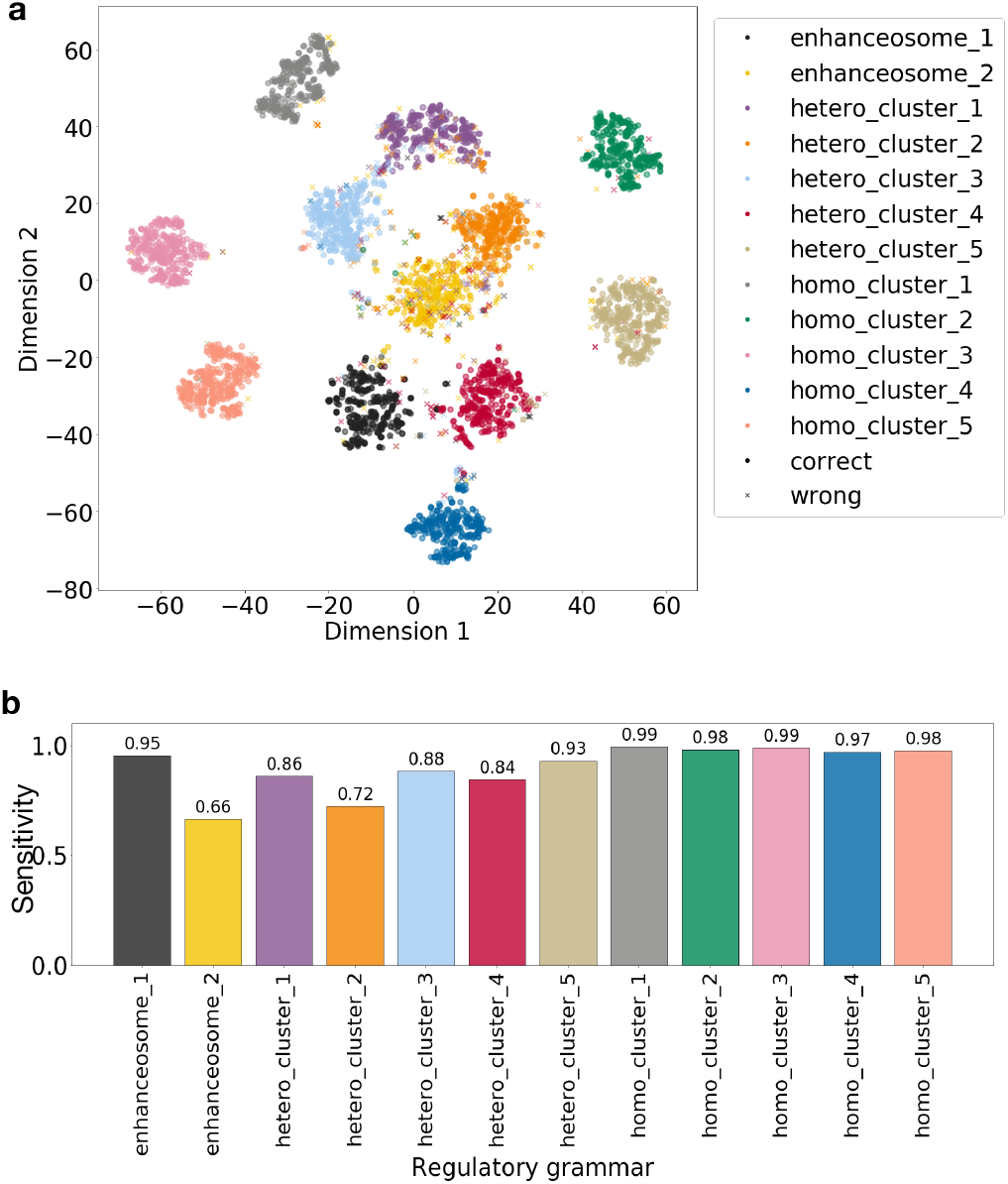
Regulatory grammar can be learned by ResNet despite heterogeneity in the regulatory sequences. (a) Actual labels of simulated regulatory grammar of the TFBS overlaid on t-SNE visualization of TFBS saliency values across neurons. (b) The sensitivity of predicted labels in (a) across regulatory grammars.

These results suggest that the model trained on regulatory sequences with heterogenous output categories can still largely capture the regulatory grammars that are essential for the heterogenous multi-label classification.

### Regulatory grammar can be learned by ResNet when a large fraction of TFBSs are not in grammars and there is heterogeneity in the regulatory sequences

In all previous prediction tasks, the simulated TFBSs in the input sequences are always in a regulatory grammar. However, in the real regulatory sequences, it is likely that only a fraction of TFBS are in regulatory grammars, while others are individual motifs scattered along the sequence. To mimic this scenario, we simulated a set of regulatory sequences with 80% of TFBSs randomly scattered in the sequence outside of any regulatory grammar and 20% of TFBSs in regulatory grammar.

We trained a ResNet model on this 80% non-grammar TFBSs dataset with the five heterogenous classes as output categories against 8-mer shuffled negatives. We found that the TFBSs outside of the regulatory grammars (single TFBS) have lower saliency values compared to the TFs in simulated regulatory grammars (Figure 5a) except for those in enhanceosome 2.

**Figure 5.**
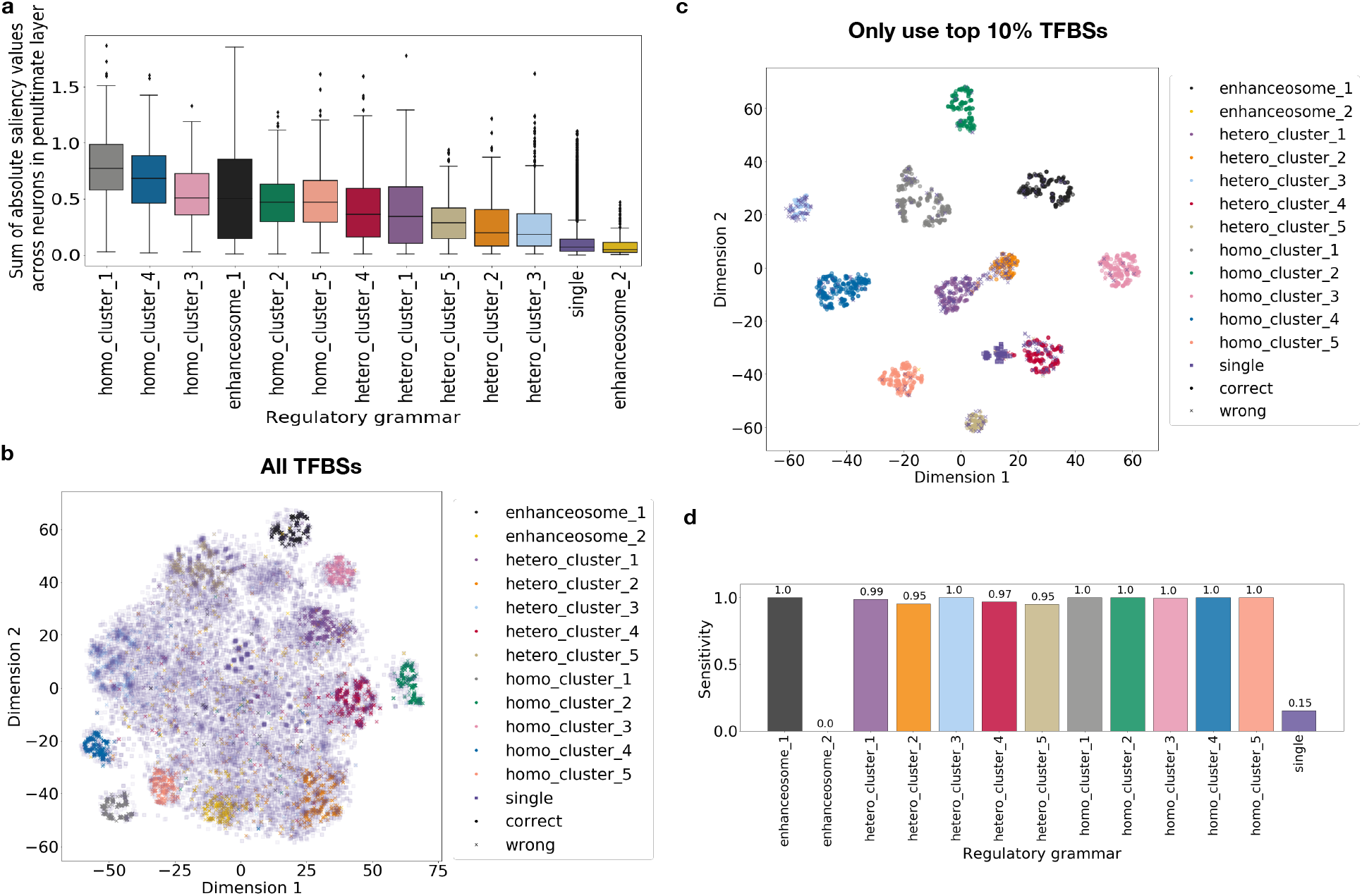
Regulatory grammar can be learned by ResNet when TFBSs are outside of regulatory grammars and there is heterogeneity in the regulatory sequence categories. (a) Sum of saliency values for TFBSs in each regulatory grammar across neurons in penultimate layer (b) Actual labels of simulated regulatory grammar of the TFBS overlaid on t-SNE visualization of TFBS saliency values across neurons. (c) Actual labels of simulated regulatory grammar of the TFBS filtered to only those in the top 10% sum of saliency values across neurons in penultimate layer overlaid on the t-SNE visualization. (d) The sensitivity of predicted labels in (c) across regulatory grammars.

Next, we performed unsupervised clustering analysis as in the previous sections (Figure 5b). Although the TFBSs in regulatory grammars still cluster, many of the TFBSs outside of regulatory grammar overlap the TFBSs in regulatory grammars in t-SNE space. This makes identifying the regulatory grammars challenging. To better reconstruct the regulatory grammar from the unsupervised clustering analysis, we took advantage of the fact that the non-grammar TFBSs have lower saliency values and only kept the TFBSs with top 10% sum of saliency values across neurons in the penultimate layer. Intuitively, this filtering helps improve the reconstruction of regulatory grammar by only focusing on TFBSs with high influence on the prediction. We repeated the unsupervised clustering analysis on these filtered TFBSs (Figure 5c). We found that nearly all TFBSs outside of regulatory grammars are filtered out (97.7%) and a smaller fraction of TFBSs in regulatory grammars are filtered (59.3%). After filtering, the remaining TFBSs are sufficient to reconstruct 11 of the 12 simulated regulatory grammars. The regulatory grammar that we failed to reconstruct, enhanceosome 2, has the lowest sum of saliency values across neurons in the penultimate layer (Figure 5a), suggesting that is was not important to learn this grammar to obtain accurate predictions. The neural network may achieve accurate predictions through elimination and therefore do not need to learn all 12 regulatory grammars.

These results suggest that even with only a small fraction of TFBSs in regulatory grammars and heterogeneity in the output categories, we can still reconstruct most of the simulated regulatory grammars.

### Regulatory grammar cannot be learned if multiple grammars are able to distinguish one regulatory sequence class from another

As shown in Figure 4 and Figure 5, some regulatory grammars, especially enhanceosome 2, are reconstructed from ResNet model with limited accuracy. This suggests that the “essentiality” of a regulatory grammar may influence the ability to reconstruct regulatory grammars from the model. In other words, if a neural network can make accurate predictions without learning certain regulatory grammars, then these non-essential regulatory grammars may not be learned during training and therefore cannot be reconstructed from the resulting model. To further investigate this hypothesis, we simulated three heterogenous regulatory classes (Table 1) with non-overlapping subsets of regulatory grammars, so that multiple regulatory grammars could distinguish one heterogenous regulatory class from another. Then we trained the model against TF-shuffled negative sequences. By setting up the training this way, the model will have to distinguish sequences with TFBSs in regulatory grammars from those with TFBSs not in regulatory grammars. However, the model does not need to learn all the regulatory grammars or distinguish one regulatory grammar from the other to make accurate predictions.

**Table 1.** Simulated heterogenous regulatory sequence classes with multiple regulatory grammars that can distinguish one class from another.

|  | *Regulatory grammar used in*<br>*the first type of sequence* | *Regulatory grammar used in*<br>*the second type of sequence* |
| --- | --- | --- |
| *Heterogeneous Regulatory*<br>*Sequence Class 1* | homotypic cluster 1,<br>homotypic cluster 2 | homotypic cluster 4,<br>heterotypic cluster 4 |
| *Heterogeneous Regulatory*<br>*Sequence Class 2* | heterotypic cluster 1,<br>heterotypic cluster 2 | homotypic cluster 5,<br>enhanceosome 1 |
| *Heterogeneous Regulatory*<br>*Sequence Class 3* | homotypic cluster 3,<br>heterotypic cluster 3 | heterotypic cluster 5,<br>enhanceosome 2 |

As expected, the model performed well at distinguishing positives and negatives (average auROC 0.995, auPR 0.978). However, when visualizing the saliency values of TFBSs of the neurons in the penultimate layer, there is limited resolution to recover individual regulatory grammars; multiple regulatory grammars have similar saliency profiles and overlap in the t-SNE space (Figure 6a). More specifically, the grammars that co-occur in the same regulatory sequence classes tend to cluster together. For example, in regulatory sequence class 1, homotypic cluster 4 clustered with homotypic cluster 1; in regulatory class 2, homotypic cluster 5 clustered with heterotypic cluster 2 and heterotypic cluster 1; in regulatory sequence class 3, homotypic cluster 3 clustered with heterotypic cluster 5. However, the remaining regulatory grammars, including homotypic cluster 2, heterotypic cluster 3, heterotypic cluster 4, enhanceosome 1, and enhanceosome 2, are scattered in the t-SNE visualization (Figure 6a). The regulatory grammars that are scattered show lower sum of saliency values across neurons, suggesting lower attention they received from the neural network (Figure 6b). This observation is consistent with our hypothesis that if there are multiple regulatory grammars that can distinguish one class of sequences from another, the neural network will not learn to distinguish one regulatory grammar from another nor learn all the distinct regulatory grammars. This scenario is likely to happen in many real enhancer classification tasks and would make reconstruction of individual regulatory grammars challenging.

**Figure 6.**
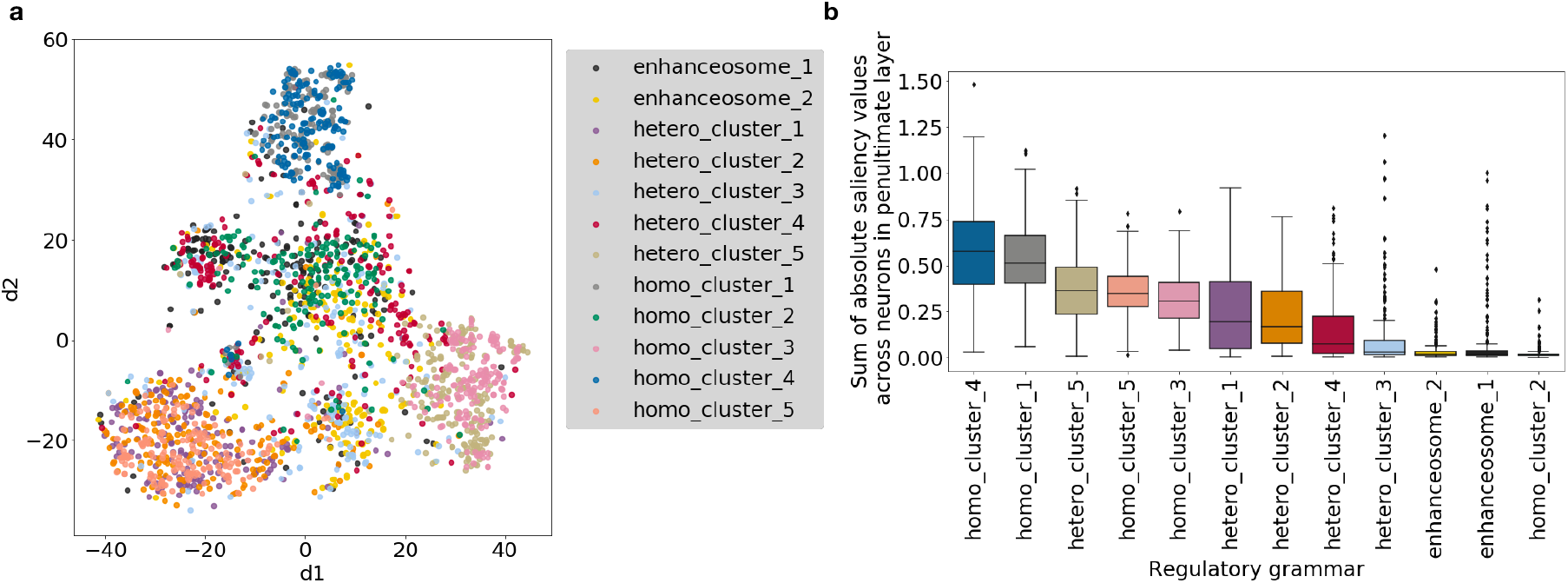
The ResNet model fails to learn the correct representation of individual grammars when there are multiple regulatory grammars that can distinguish one heterogenous regulatory class from another. Actual labels of simulated regulatory grammar of the TF binding sites overlaid on t-SNE visualization of TFBS saliency values across neurons. b) Sum of saliency values for TFBSs in each regulatory grammar across neurons in the penultimate layer.

## Discussion

We trained a variant of DNNs, ResNets, to model sequences with simulated regulatory grammars (combinatorial binding of TFs). Then we developed a gradient-based unsupervised clustering approach to interpret the features learned by neurons in the intermediate layers of the neural network. We found that ResNets can model the simulated regulatory grammars even when there is heterogeneity in the regulatory sequences and a large fraction of TFBSs outside of regulatory grammars.

We also identified scenarios when the ResNet model failed to learn the regulatory grammar. The networks strive to learn simple representations of the training data. As a result, the ResNet models in our studies failed to learn the simulated regulatory grammar when there is a lack of constraint in negative training samples or between the positive output categories. For instance, we found that the choice of negative training samples influences the ability of the neural network to learn regulatory grammar. The model trained against no negatives or short k-mer shuffled negatives (k=1, 2, 4, 6) or very long k-mer shuffled negatives (k=12) did not learn accurate representation of the regulatory grammar and often misclassified TF-shuffled negatives as positives. The model trained against 8-mer shuffled negatives performed the best when evaluated on the TF-shuffled negatives. This is because when shorter k-mers (k<=6) are used to generate the negative training samples, the neural network can distinguish the positives from negatives by learning the individual TF motifs, many of which are longer than 6 bp, rather than learning the regulatory grammar of the TFs. With longer k-mers (k=12), the reason is likely that k-mers are not well shuffled in the negatives and very similar to the positives. Indeed, the ResNet model trained against 12-mer shuffled negatives has a low accuracy (auPR 0.506). The 8-mer shuffled negative provides a sweet spot where the negatives are well shuffled and the network is forced to learn the TF motifs and regulatory grammars. Another challenging situation occurs when there are multiple sequence features that can distinguish one output category from another. Under this scenario, it is not necessary for the neural network to accurately learn all the features nor distinguish one feature from another.

In addition to these scenarios, there is also another situation in which the ResNet model failed to learn the regulatory grammar. When the majority of the TFBSs are not in a regulatory grammar, the non-grammar TFBSs overlap those in regulatory grammars in the unsupervised clustering analysis and make it impossible to recover the grammars. Fortunately, we could use the observation that many of the TFBSs outside of regulatory grammars have low saliency values to filter out those TFBSs, and focus the unsupervised clustering analysis on TFBS with high saliency values to improve the accuracy of grammar reconstruction. This gradient magnitude-based filtering method may be less efficient when there is an overwhelmingly large number of TFBSs outside of regulatory grammar and larger sample sizes might be needed to train the neural network to better retrieve the regulatory grammars.

While we demonstrate potential to interpret biologically relevant patterns learned by deep neural network models in some realistic scenarios, our work has several caveats. First, the synthetic dataset and proposed methods assume that combinatorial binding of TFs does not change their motifs. However, this assumption may not be always true. In vitro analyses of the combinatorial binding of pairs of TFs indicate that many pairs of TFs have different binding motifs when they bind together compared to their consensus motifs (Jolma et al., 2015). Although there is nothing preventing the neural network from learning such altered motifs, the unsupervised clustering methods based on individual TFBS may have limited accuracy in identifying such altered motifs. Second, we did not simulate noisy labels in the synthetic dataset which could occur in the real regulatory sequence prediction tasks. The common methods of experimentally finding enhancers, such as ChIP-seq on histone modifications, DNase-Seq, CAGE-seq, and MPRAs, often produce mislabeled regulatory regions and vague region boundaries. This could be improved in the future by integrating methods for learning from noisy labeled data.

In summary, we demonstrated the power and limitations of deep convolutional neural networks at modeling regulatory grammars and provided a backpropagation gradient based unsupervised learning approach to retrieve and interpret the patterns learned by inner layers of the neural network. Our work indicates that DNNs can learn biologically relevant TFBS combinations in certain settings with carefully defined training data; however, in many common scenarios, we should be cautious when interpreting the biological implications of features learned by DNA-sequence-based DNNs.

## Materials and Methods

### Simulated sequence generation and analysis

#### Simulation of regulatory grammar

We used TF binding motifs from the HOCOMOCO v11 database (Kulakovskiy et al., 2017). To make sure that the TF motifs are distinct and diverse, we select one TF from each TF subfamily. This results in a set of 26 TFs (Supplementary Table 1). Then the selected TFs are arranged into three types of regulatory grammar representing homotypic clusters, heterotypic clusters, and enhanceosomes.

For the homotypic cluster, we simulated multiple non-overlapping occurrences (3-5) of the same TF in a small window (120 bp) at random locations. For the heterotypic clusters, we simulated a set of four diverse TFs in a small window (120 bp) at random non-overlapping locations. Each TF occurs once in the heterotypic cluster. For the enhanceosome, we simulated a set of four TFs in a small window with fixed order and spacing. Because it is possible in real enhancers that the same TF factor is used in different regulatory grammars, we allow some of TFs to occur in more than one grammar. We simulated five homotypic TF clusters, five heterotypic clusters and two enhanceosomes (Supplementary Table 2).

#### Simulation of regulatory sequences with different regulatory grammars

To mimic common enhancer prediction tasks, such as predicting enhancers from different cellular contexts, we designed twelve regulatory sequence classes (Supplementary Table 3) with each regulatory sequence class representing one type of enhancer (e.g., enhancers active in a given context). Sequences in each class have two different regulatory grammars. Because it is possible that the same regulatory grammar is used in regulatory sequences in different cellular contexts, we allow one regulatory grammar occur in two different regulatory sequence classes. For instance, the first regulatory sequence class has homotypic cluster 1 and heterotypic cluster 1, then the second regulatory sequence class has heterotypic cluster 1 and homotypic cluster 2 and then the third regulatory sequence class has homotypic cluster 2 and heterotypic cluster 3, etc. Next, we randomly generated background DNA sequences of 3000 bp based on equal probability of A, G, C, T and inserted 2-4 of each simulated regulatory grammar at random locations into these background sequences based on the corresponding regulatory class.

#### Multiclass classification and heterogenous class classification

We performed two types of classification: i) multiclass classification in which each output neuron represents a homogenous set of regulatory sequences and ii) heterogenous class classification in which each output neuron represents a heterogenous set of regulatory sequences. The heterogenous class classification task assumes that in the real enhancer prediction tasks,enhancers in one category (e.g., specific cellular context) may have a heterogenous set of sequences harboring different sets of regulatory grammars.

The multiclass classification task has twelve homogeneous output classes, each one corresponding to sequences representing one regulatory sequence class. The heterogenous class classification (Supplementary Table 4) has five heterogeneous output classes, each one corresponding to a subset of regulatory sequence classes. More specifically, heterogeneous class 1 has regulatory sequence class 1, 3, and 5; heterogeneous class 2 has regulatory sequence class 2, 4, and 6; heterogeneous class 3 has regulatory sequence class 7, 9, and 11; heterogeneous class 4 has regulatory sequence class 5, 8, and 10; heterogeneous class 2 has regulatory sequence class 1, 6, and 12.

#### Negative sequences

We used three approaches to generate negatives when training the classifiers: no negatives, k-mer shuffled negatives, and TF-shuffled negatives. For the k-mer shuffled negative sequence set, we matched the frequency of k-mers (k=1, 2, 4, 8, 12) in the negatives to the simulated regulatory (positive) sequences. For the TF-shuffled sequence set, we shuffled the TFBS of the simulated regulatory sequences.

### Model design and training

DNNs have achieved the state-of-art performance on regulatory sequence prediction (Quang and Xie, 2016; Zhou and Troyanskaya, 2015). The integration of a convolution operation into standard neural networks enables learning common patterns that occur at different spatial positions, such as TF motifs in the DNA sequences. Here we use a residual deep convolutional neural networks (ResNets), a variant of DNNs that allows connections between non-sequential layers (He et al., 2016) to model the regulatory sequences. Each simulated DNA sequence is one-hot-encoded, which is represented by a sequence length × 4 matrix with columns representing A, G, C and T.

The basic layers in the network include a convolutional layer, batch normalization layer, pooling layer, and fully connected layer. Every two convolutional layers are grouped into a residual block where an identity shortcut connection adds the input to the residual block to the output of the residual block. This additional identity mapping is an efficient way to deal with vanished gradients that occur in neural networks with large depth and improves performance in many scenarios. The batch normalization layers are added after the activation of each residual block. Batch normalization (Ioffe and Szegedy, 2015) helps reduce the covariance shift of the hidden unit and allows each layer of a neural network to learn more independently of other layers. The pooling layers are added after each batch normalization layer. Finally, a dense (fully connected) layer and an output layer are added at the top of the neural network. We used 4 residual blocks, each has two convolutional layers with 32 neurons. The final residual block is connected to a dense layer with 32 neurons and then connected to output layer (Supplementary Figure 1). We found the above neural network structure (ResNet) performed well in all of our simulation tasks while a 3-layer convolutional neural network with alternating convolutional layers and maxpooling layers cannot, suggesting the benefit of using a much deeper neural network at modeling enhancer regulatory grammar.

We used rectified (ReLU) activation for all the residual blocks and sigmoid activation for the output fully connected layer activation. We used binary cross-entropy as the loss function and Adam (Kingma and Ba, 2014) as the optimizer.

### Model interpretation and grammar reconstruction

#### Computing saliency values with respect to neurons

We considered two gradient calculating approaches for estimating the importance of each nucleotide in the input sequence with respect to each neuron’s activation. The first is guided back-propagation in which we calculated the gradient of the neuron of interest with respect to the input through guided back-propagation and then multiplied the gradient by input sequences. The second is calculating the DeepLIFT score (Shrikumar et al., 2017) of the neuron of interest with respect to the input using the DeepLIFT algorithm implemented in SHAP (Lundberg and Lee, 2017) against the TF-shuffled negatives and then multiplying the DeepLIFT score by input sequences. We refer the resulting values from the above as saliency values and the vector of saliency values for an input sequence as saliency map. We found that the saliency maps calculated using DeepLIFT approach performed the better than guided back-propagation (Supplementary Figure 2). Therefore, for all the main text results we present were calculated with the DeepLIFT approach.

#### Analysis of TF saliency maps

To analyze which TFs are learned by a specific neuron, we calculate the gradient of a TF binding site with respect to a neuron by averaging a 10 bp window from the start position of the TF binding site. Then, we visualize the distribution of saliency values of the binding sites of each TF in a specific regulatory grammar with respect to a neuron with box plot.

The median saliency values of the binding sites of each TF in a specific regulatory grammar with respect to neurons is stored in a matrix with the shape of number of neurons by the number of TFs. This data matrix is first scaled by column to identify which neurons mostly detect the TF and the scaled matrix is used to generate a heatmap. Then, we performed hierarchical clustering with k=12 (12 is the number of simulated regulatory grammars) or k=13 (when there are non-grammar TFBSs) for both neurons and TFs based on the same data matrix.

#### t-SNE and k-means clustering of TFBS

To reconstruct the regulatory grammar and evaluate how accurately neurons in a layer capture the simulated regulatory grammar, we performed a two-dimensional t-SNE and a k-means clustering (k=12) of TFBS using their saliency value profiles across neurons in a layer. To assign the name of regulatory grammar of a predicted cluster, we used a majority vote, which is the majority of the true labels of regulatory grammar in that cluster. We visualize the k-means clustering by overlaying the predicted regulatory grammar from k-means clustering on top of the t-SNE visualization. We evaluated the performance at reconstructing the regulatory grammar by two metrics: the accuracy ((TP+TN)/All) and the sensitivity (TP/(TP+FN)) of the regulatory grammar predictions.

## Acknowledgements

We are grateful to Dennis Kostka, David Rinker, Laura Colbran, and members of the Capra Lab for helpful discussions and comments on the manuscript. This work was supported by the National Institutes of Health (USA) awards R35GM127087 and R01GM115836 (JAC) and the Burroughs Wellcome Fund (JAC).

