## Supplementary Figure for "LEARNING AND INTERPRETING THE GENE REGULATORY GRAMMAR IN A DEEP LEARNING FRAMEWORK"

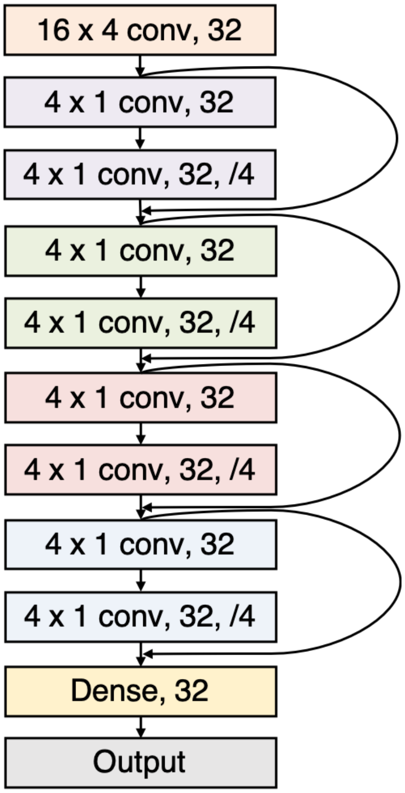


Figure 1: The structure of the ResNet model.

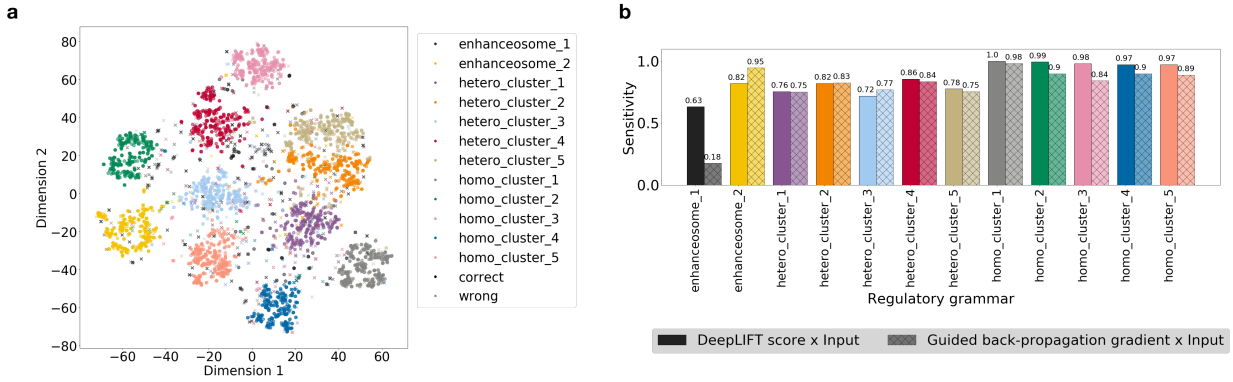


Figure 2: The DeepLIFT score is better at reconstructing the regulatory grammar compared to guided back-propagation gradient. a. True and predicted labels of simulated regulatory grammar of the TF binding sites overlaid on t-SNE visualization. b. The sensitivity (TP/TP+FN) of predicted labels of regulatory grammar using DeepLIFT score x Input or Guided back-propagation gradient

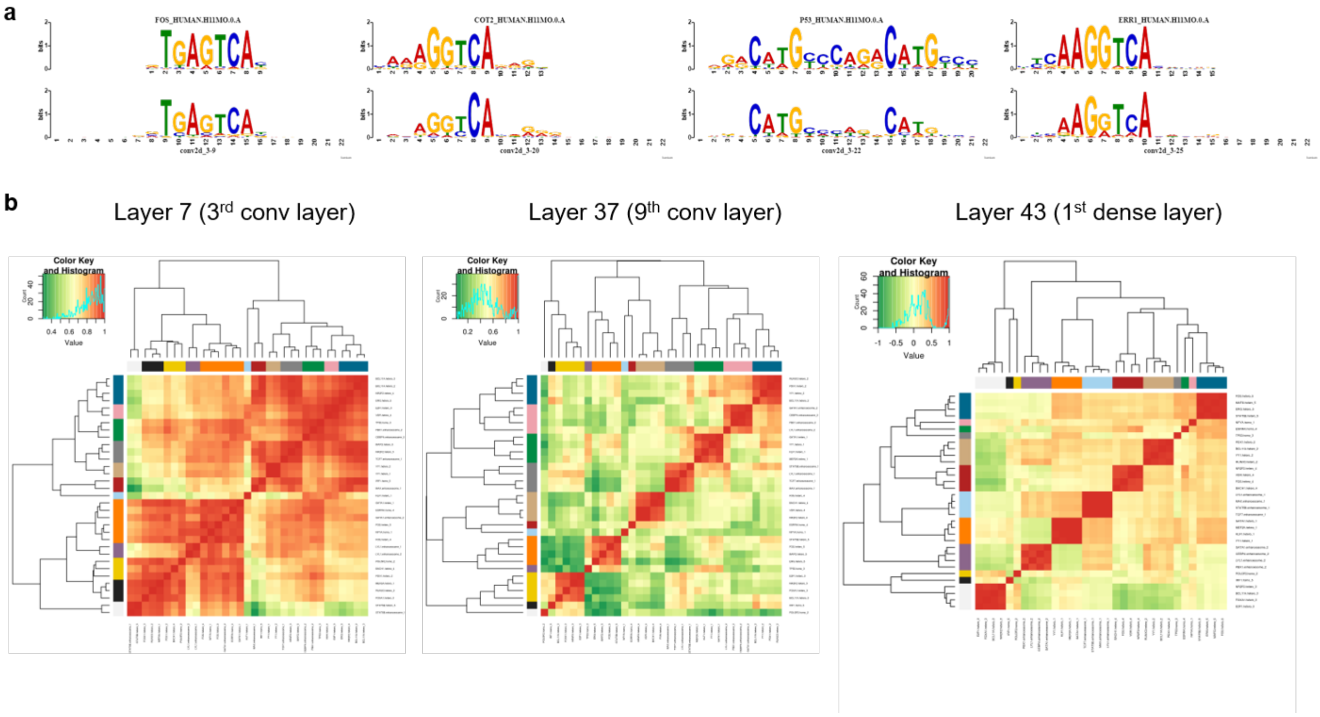


Figure 3: The neural network learned individual TF binding motifs in the lower convolutional layer and gradually build up its understanding of regulatory grammar in higher level layers. a. Simulated TF motifs are learned by neurons in the third convolutional layer. From left to right are four selected examples, neuron 9 learned the FOS motif; neuron 20 learned the COT2 motif; neuron 22 learned the P53 motif; neuron 25 learned ERR1 motif. b. From layer 7 (third convolutional layer) to Layer 43 (the penultimate dense layer), the ResNet model gradually learned the regulatory grammar. The correlation matrix of the saliency value profiles of TFs in a specific regulatory grammar is plotted as the heatmap. In layer 7, TFs from the same regulatory grammar are not clustered. In layer 37, TFs within the same regulatory grammar begin to have a higher correlation. In layer 43, TFs within the same regulatory grammar have near perfect correlation.

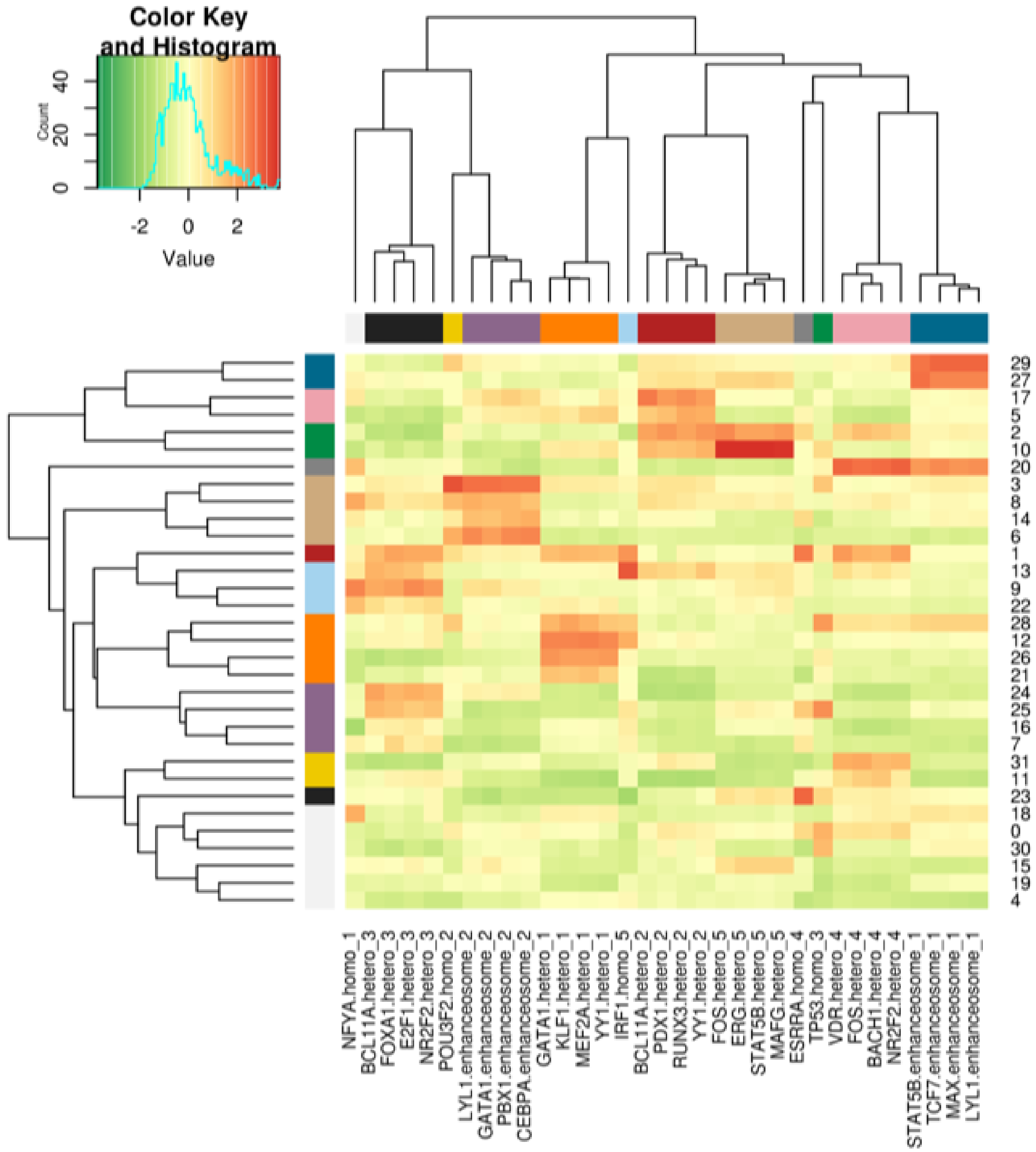


Figure 4: ResNet model trained on simulated regulatory sequences and 8-mer shuffled negatives. Heatmap of the median gradient of the binding sites of each TF in a specific regulatory grammar (x axis) across neurons of the penultimate layer (y axis). The order of x and y axis labels are determined by hierarchical clustering shown on side. The color bars indicate the group label assigned by hierarchical clustering.

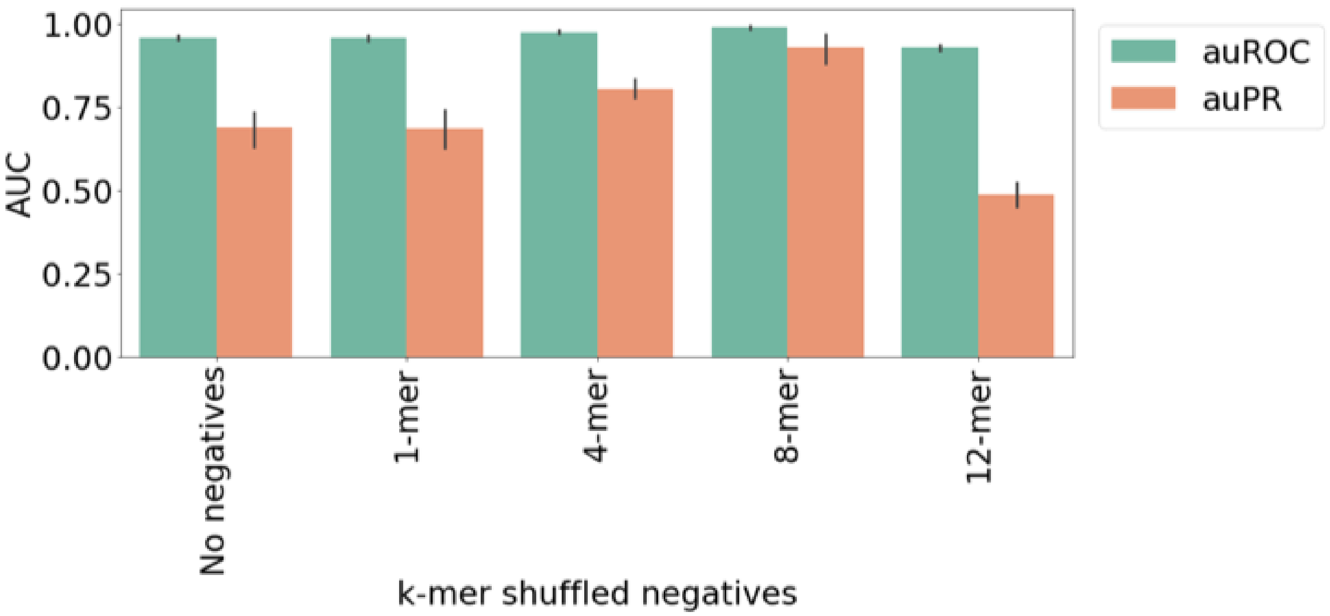


Figure 5: ResNet model accuracy trained for heterogeneous multilabel classification with no negatives or against k-mer shuffled negatives.
