## Supplementary Table for "LEARNING AND INTERPRETING THE GENE REGULATORY GRAMMAR IN A DEEP LEARNING FRAMEWORK"

Table 1: Transcription factor used in constructing regulatory grammars

| TF | logo | regulatory rammar |
|---|---|---|
| NFYA | 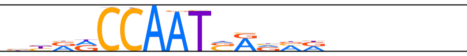 | homo_cluster_1 |
| POU3F2 | 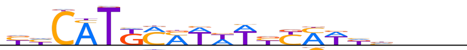 | homo_cluster_2 |
| TP53 | 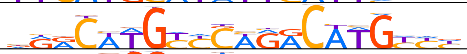 | homo_cluster_3 |
| ESRRA | 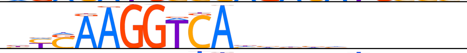 | homo_cluster_4 |
| IRF1 | 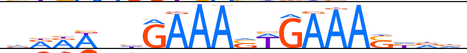 | homo_cluster_5 |
| KLF1 | 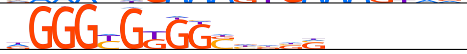 | hetero_cluster_1 |
| MEF2A | 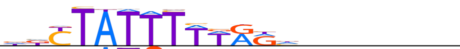 | hetero_cluster_1 |
| YY1 | 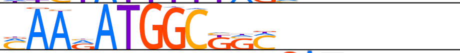 | hetero_cluster_1, hetero_cluster_2 |
| GATA1 | 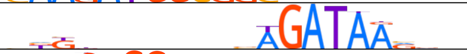 | hetero_cluster_1, enhanceosome_2 |
| RUNX3 | 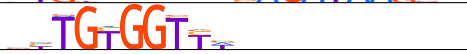 | hetero_cluster_2 |
| PDX1 | 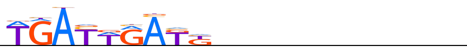 | hetero_cluster_2 |
| BCL11A | 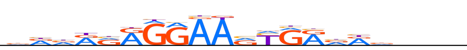 | hetero_cluster_2, hetero_cluster_3 |
| E2F1 | 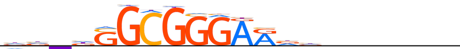 | hetero_cluster_3 |
| FOXA1 | 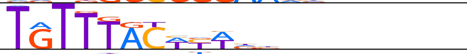 | hetero_cluster_3 |
| NR2F2 | 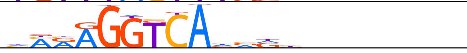 | hetero_cluster_3, hetero_cluster_4 |
| VDR | 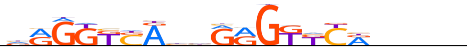 | hetero_cluster_4 |
| BACH1 | 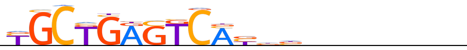 | hetero_cluster_4 |
| FOS | 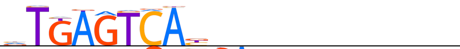 | hetero_cluster_4, hetero_cluster_5 |
| MAFG | 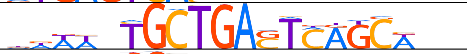 | hetero_cluster_5 |
| ERG | 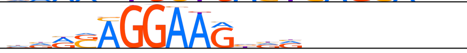 | hetero_cluster_5 |
| STAT5B | 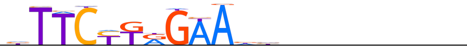 | hetero_cluster_5, enhanceosome_1 |
| TCF7 | 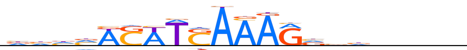 | enhanceosome_1 |
| MAX | 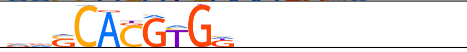 | enhanceosome_1 |
| LYL1 | 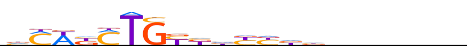 | enhanceosome_1, enhanceosome_2 |
| PBX1 | 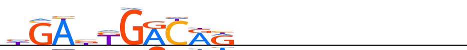 | enhanceosome_2 |
| CEBPA | 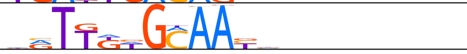 | enhanceosome_2 |

Table 2: Simulated regulatory grammar

| type | name | TFs |
|---|---|---|
| homotypic cluster | homo_cluster_1 | NFYA |
| homotypic cluster | homo_cluster_2 | POU3F2 |
| homotypic cluster | homo_cluster_3 | TP53 |
| homotypic cluster | homo_cluster_4 | ESRRA |
| homotypic cluster | homo_cluster_5 | IRF1 |
| heterotypic cluster | hetero_cluster_1 | KLF1, MEF2A, YY1, GATA1 |
| heterotypic cluster | hetero_cluster_2 | RUNX3, PDX1, BCL11A, YY1 |
| heterotypic cluster | hetero_cluster_3 | E2F1, FOXA, NR2F2, BCL11A |
| heterotypic cluster | hetero_cluster_4 | VDR, BACH1, FOS, NR2F2 |
| heterotypic cluster | hetero_cluster_5 | MAFG, ERG, STAT5B, FOS |
| enhanceosome | enhanceosome_1 | TCF7, MAX, LYL1, STAT5B |
| enhanceosome | enhanceosome_2 | PBX1, CEBPA, GATA1, LYL1 |

Table 3: Simulated regulatory classes

| regulatory sequence class | regulatory grammars |
|---|---|
| regulatory_class1 | homo_cluster_1, hetero_cluster_1 |
| regulatory_class2 | hetero_cluster_1, homo_cluster_2 |
| regulatory_class3 | homo_cluster_2, hetero_cluster_2 |
| regulatory_class4 | hetero_cluster_2, homo_cluster_3 |
| regulatory_class5 | homo_cluster_3, hetero_cluster_3 |
| regulatory_class6 | hetero_cluster_3, homo_cluster_4 |
| regulatory_class7 | homo_cluster_4, hetero_cluster_4 |
| regulatory_class8 | hetero_cluster_4, homo_cluster_5 |
| regulatory_class9 | homo_cluster_5, hetero_cluster_4 |
| regulatory_class10 | hetero_cluster_5, enhanceosome_1 |
| regulatory_class11 | enhanceosom_1, enhanceosome_2 |
| regulatory_class12 | enhanceosom_2, homo_cluster_1 |

Table 4: Simulated heterogenous regulatory classes

| heterogenous regulatory sequence class | regulatory classes |
|---|---|
| heterogenous_regulatory_class1 | regulatory_class1, regulatory_class3, regulatory_class5 |
| heterogenous_regulatory_class2 | regulatory_class2, regulatory_class4, regulatory_class6 |
| heterogenous_regulatory_class3 | regulatory_class7, regulatory_class9, regulatory_class11 |
| heterogenous_regulatory_class4 | regulatory_class5, regulatory_class8, regulatory_class10 |
| heterogenous_regulatory_class5 | regulatory_class1, regulatory_class6, regulatory_class12 |

Table 5: Simulated no-overlap heterogenous regulatory classes

| heterogenous regulatory sequence class | regulatory classes | regulatory grammars |
|---|---|---|
| heterogenous_regulatory_class1 | regulatory_class1, regulatory_class4, regulatory_class7, regulatory_class10 | homo cluster 1, hetero cluster 1, hetero cluster 2, homo cluster 3, homo cluster 4, hetero cluster 4, hetero cluster 5, enhanceosome 1 |
| heterogenous_regulatory_class2 | regulatory_class2, regulatory_class5, regulatory_class9, regulatory_class11 | hetero cluster 1, homo cluster 2, homo cluster 3, hetero cluster 3, homo cluster 5, hetero cluster 4, enhanceosome 1, enhanceosome 2 |
| heterogenous_regulatory_class3 | regulatory_class3, regulatory_class6, regulatory_class10, regulatory_class12 | homo cluster 2, hetero cluster 2, hetero cluster 3, homo cluster 4, hetero cluster 5, enhanceosome 1, enhanceosom 2, homo cluster 1 |
